# Coordinated modulation of multiple processes through phase variation of a c-di-GMP phosphodiesterase in *Clostridioides difficile*

**DOI:** 10.1101/2021.10.30.466581

**Authors:** Leila M. Reyes Ruiz, Kathleen A. King, Elizabeth M. Garrett, Rita Tamayo

**Affiliations:** Department of Microbiology and Immunology, University of North Carolina School of Medicine, Chapel Hill, North Carolina, USA

**Author notes:** Corresponding Author Marsico Hall Rm 6101, 125 Mason Farm Rd, CB#7290, Chapel Hill, NC 27599-7280.

**Keywords:** cyclic diguanylate, clostridium, biofilm, motility, toxin, virulence

## Abstract

The opportunistic nosocomial pathogen *Clostridioides difficile* exhibits phenotypic heterogeneity through phase variation, a stochastic, reversible process that modulates expression. In *C. difficile*, multiple sequences in the genome undergo inversion through site-specific recombination. Two such loci lie upstream of *pdcB* and *pdcC*, which encode phosphodiesterases (PDEs) that degrade the signaling molecule c-di-GMP. Numerous phenotypes are influenced by c-di-GMP in *C. difficile* including cell and colony morphology, motility, colonization, and virulence. In this study, we aimed to assess whether PdcB phase varies, identify the mechanism of regulation, and determine the effects on intracellular c-di-GMP levels and regulated phenotypes. We found that expression of *pdcB* is heterogeneous and the orientation of the invertible sequence, or ‘*pdcB* switch’, determines expression. The *pdcB* switch contains a promoter that when properly oriented promotes *pdcB* expression. Expression is augmented by an additional promoter upstream of the *pdcB* switch. Mutation of nucleotides at the site of recombination resulted in phase-locked strains with significant differences in *pdcB* expression. Characterization of these mutants showed that the *pdcB* locked-ON mutant has reduced intracellular c-di-GMP compared to the locked-OFF mutant, consistent with increased and decreased PdcB activity, respectively. These alterations in c-di-GMP had concomitant effects on multiple known c-di-GMP regulated processes. These results indicate that phase variation of PdcB allows *C. difficile* to coordinately diversify multiple phenotypes in the population to enhance survival.

## INTRODUCTION

*Clostridioides difficile* is a gram-positive, spore-forming, obligate anaerobe that causes antibiotic-associated intestinal disease ranging from mild diarrhea to pseudomembranous colitis in susceptible hosts. *C. difficile* is one of the leading causes of hospital acquired infections in the United States and has been identified as an urgent threat by the Center for Disease Control and Prevention (CDC) (1, 2). Pathogenesis of *C. difficile* infection (CDI) is largely mediated by the secreted toxins TcdA and TcdB (3-5), which inactivate Rho-family GTPases (6, 7). Inactivation of Rho GTPases of intestinal epithelial cells leads to dissociation of the actin cytoskeleton at the epithelial barrier, disruption of tight junctions, and cell death, which results in inflammation and diarrheal symptoms (8, 9).

Several recent studies have shown that *C. difficile* exhibits heterogeneity in gene expression and associated phenotypes (10-13). Phase variation is one way bacteria introduce phenotypic heterogeneity into a clonal population, helping ensure the survival of the population if exposed to adverse conditions (14, 15). Typically, phase variation modulates the production of surface factors such as flagella, fimbriae, and exopolysaccharides; in host-associated bacteria, the process can influence immune system evasion, colonization, and virulence (14, 16). Several genetic and epigenetic mechanisms can drive phase variation, including conservative site-specific DNA recombination in which a serine or tyrosine recombinase mediates the reversible inversion of a DNA sequence (17, 18). The invertible sequence, or “switch”, is flanked by short, inverted repeats and contains the information necessary to regulate the expression of the adjacent gene or operon.

In the *C. difficile* epidemic-associated strain R20291, seven sequences undergo inversion (10, 11, 19), and three have been shown to mediate phase variation. These switches modulate expression of: *cwpV*, which encodes a cell wall protein involved in bacteriophage resistance (10, 20); *cmrRST*, which encodes an atypical signal transduction system that impacts cell and colony morphology, surface motility, flagellar motility, and virulence (12, 19); and the *flgB* operon, a large operon encoding flagellar hook and basal body components as well as the sigma factor SigD (11, 21, 22). SigD coordinates expression of additional flagellar operons and positively regulates expression of the *tcdA* and *tcdB* toxin genes (23, 24). Accordingly, toxin production is indirectly subject to phase variation via the flagellar switch (11, 21). Phase variation via the *cwpV* and flagellar switches occur through RNA-mediated mechanisms post-transcription initiation, but the gene regulatory mechanisms of the remaining switches remain unknown (10, 11, 22).

The signaling molecule cyclic diguanylate monophosphate (c-di-GMP) regulates the transition between motile and sessile lifestyles in numerous bacterial species (25, 26). c-di-GMP is synthesized by diguanylate cyclases (DGCs) containing a GGDEF domain and degraded by phosphodiesterases (PDEs) containing EAL or HD-GYP domains (26). These enzymes are controlled at the transcriptional and post-translational levels, and their opposing activities control the intracellular c-di-GMP concentration. Like many bacterial species, *C. difficile* encodes dozens of c-di-GMP metabolic enzymes; the ribotype 027 strain used in this study, R20291, encodes 14 known or putative DGCs and 17 known or putative PDEs (27, 28). In *C. difficile*, regulation of gene expression by c-di-GMP is largely mediated by riboswitches that either increase or decrease gene expression when c-di-GMP is bound (29-32). The majority of genes directly regulated via a c-di-GMP riboswitch in *C. difficile* encode surface-associated factors that impact adhesion or motility. As in other bacterial species, c-di-GMP has broad regulatory effects in *C. difficile*, inhibiting flagellar gene expression, swimming motility and toxin production (33, 34), and increasing expression of adhesin genes and associated surface behaviors such as biofilm formation (35-40).

Phase variation and c-di-GMP signaling appear to be linked in *C. difficile*. Expression of the *flgB* and *cmrRST* operons is regulated by both c-di-GMP and phase variation (31, 33, 41). In addition, two of the genes encoding proteins with tandem GGDEF and EAL domains, CDR20291_0685 (*pdcB*) and CDR20291_1514 (*pdcC*), are preceded by invertible elements and are thus likely subject to phase variation (19, 42). Overexpression of the orthologous genes from *C. difficile* 630 in *Vibrio cholerae* and *Bacillus subtilis*, which have well-described responses to c-di-GMP, are consistent with PDE function, and a *pdcB* orthologue (CD630_0757) exhibited c-di-GMP hydrolytic activity *in vitro* (27, 43). Phase variation of these PDEs is therefore poised to modulate one or more c-di-GMP-regulated phenotype. However, the impact of these PDEs on c-di-GMP and phenotypic outcomes in *C. difficile* is unknown.

In this study, we aimed to determine whether PdcB is subject to phase variation and the potential consequence to c-di-GMP signaling and *C. difficile* physiology and behavior. We found that the upstream invertible element, the “*pdcB* switch”, can be found in both orientations in multiple *C. difficile* strains. We demonstrated that *pdcB* is expressed heterogeneously in *C. difficile*, identified the switch orientation that promotes *pdcB* expression, and determined the mechanism by which the switch regulates expression. By mutating key nucleotides in the right inverted repeat, we created *pdcB* phase-locked strains that are incapable of switch inversion. These mutants differed significantly in *pdcB* expression. Using a riboswitch-based fluorescent reporter, we found that the phase-locked ON mutant had significantly lower c-di-GMP compared to the phase-locked OFF mutant, supporting PDE activity for PdcB. Consistent with these differences in c-di-GMP, the phase-locked ON mutant showed reduced swimming motility, but increased surface motility and biofilm formation that the phase-locked OFF mutant. Throughout this study, we accounted for the potential contributions of the multiple other phase variable factors, particularly PdcC and others that might influence motility and biofilm phenotypes. Together these results indicate that *C. difficile* employs phase variation of a c-di-GMP PDE to coordinately modulate multiple mechanisms of adaptation to environmental stimuli.

## RESULTS

### The orientation of the *pdcB* switch is heterogeneous in multiple *C. difficile* strains

The *pdcB* gene encodes a c-di-GMP phosphodiesterase (PDE) containing a PAS sensory domain, a degenerate GGDEF domain, and an EAL domain (Fig 1A) (27). In *C. difficile* R20291, an epidemic-associated ribotype 027 strain, the invertible element Cdi2 is 174 bp long, flanked by inverted repeats, and located 840 bp upstream of the *pdcB* coding sequence (Fig 1A)(19, 44). Sekulovic et al. previously showed that the Cdi2 sequence can be found in both orientations in R20291, indicating that the sequence undergoes inversion (19). To determine whether the Cdi2 sequence is similarly heterogeneous in other *C. difficile* strains, we employed a previously described PCR strategy, orientation-specific PCR (OS-PCR), which uses primer sets that allow the differential amplification of either Cdi2 sequence orientation, qualitatively showing the presence or absence of each (10, 11, 45). We selected four additional *C. difficile* strains representing multiple ribotypes (RT): UK1 (RT027), 630 (RT012), VPI10463 (RT087), and ATCC43598 (RT017). ATCC BAA1875 (RT078), which lacks an orthologous *pdcB* locus including the region containing the *pdcB* switch, was included as a negative control. The Cdi2 sequence was detected in both orientations in all but the BAA1875 negative control (Fig. 1B). This heterogeneity in the orientation of the Cdi2 sequence, herein named the “*pdcB* switch”, indicates that the sequence undergoes inversion in multiple *C. difficile* strains when grown in standard laboratory conditions.

**Figure 1.**
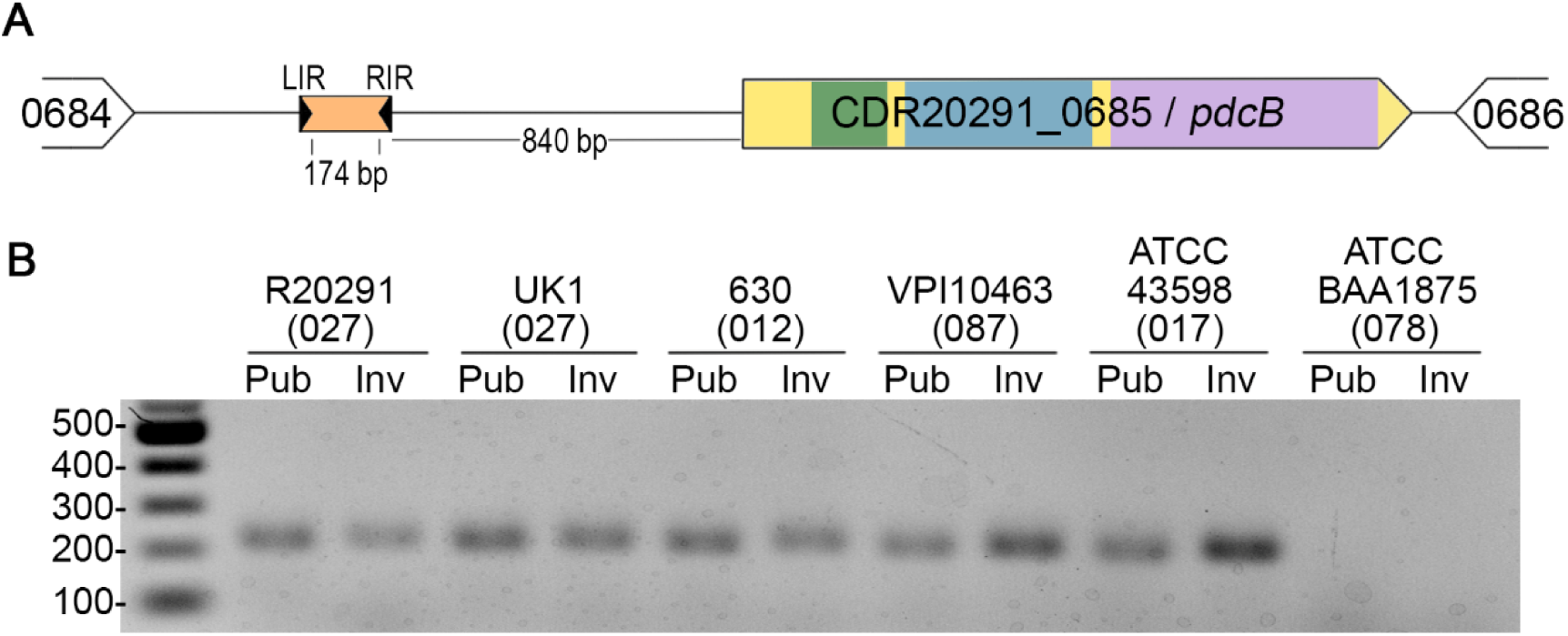
The *pdcB* switch (Cdi2) is invertible in *C. difficile*. (A) Diagram of the *pdcB* locus indicating the relative positions of the upstream invertible element (orange) and inverted repeats (black arrows). LIR – left inverted repeat, RIR – right inverted repeat. PdcB contains a PAS domain (green), a GGDEF domain (blue), and an EAL domain (purple). (B) Orientation-specific PCR for the *pdcB* switch in six different *C. difficile* strains (R20291, UK1, 630, VPI10463, ATCC43598, and ATCC BAA1875). The previously determined ribotype of each strain is indicated in parenthesis. Pub – primers designed to amplify from the sequence orientation present in the published R20291 genome sequence (Accession No. FN545816), Inv – primers designed to amplify the inverted sequence orientation.

### *pdcB* switch orientation modulates *pdcB* expression in an ON/OFF manner

To quantitatively assess the heterogeneity in *pdcB* switch orientation and its impact on *pdcB* expression, we isolated six independent colonies of R20291 grown on BHIS-agar and collected paired samples for genomic DNA and RNA isolation. The genomic DNA was subjected to quantitative PCR using the orientation-specific primers described above (OS-qPCR) to determine the percentage of a population with the switch in the orientation present in the R20291 reference genome (accession number FN545816). The abundance of *pdcB* transcript in the matched RNA sample was evaluated by quantitative reverse transcriptase PCR (qRT-PCR) with normalization to the abundance of *rpoC* transcript.

Of the six isolates, four contained the *pdcB* switch predominantly in the reference genome orientation (99.27% ± 1.21% reference) (Fig 2B, WT triangles). These isolates showed low levels of *pdcB* mRNA, 0.8% of *rpoC* transcript levels (Fig 2C, WT triangles). The remaining two isolates had the *pdcB* switch in the inverse orientation (0.38% ± 0.42% reference) (Fig 2B, WT squares). These two isolates had significantly higher *pdcB* transcript levels, 16% of *rpoC* transcript levels (Fig 2C, WT triangles). These results show that *pdcB* switch orientation and expression is bimodal in wild-type *C. difficile*.

**Figure 2.**
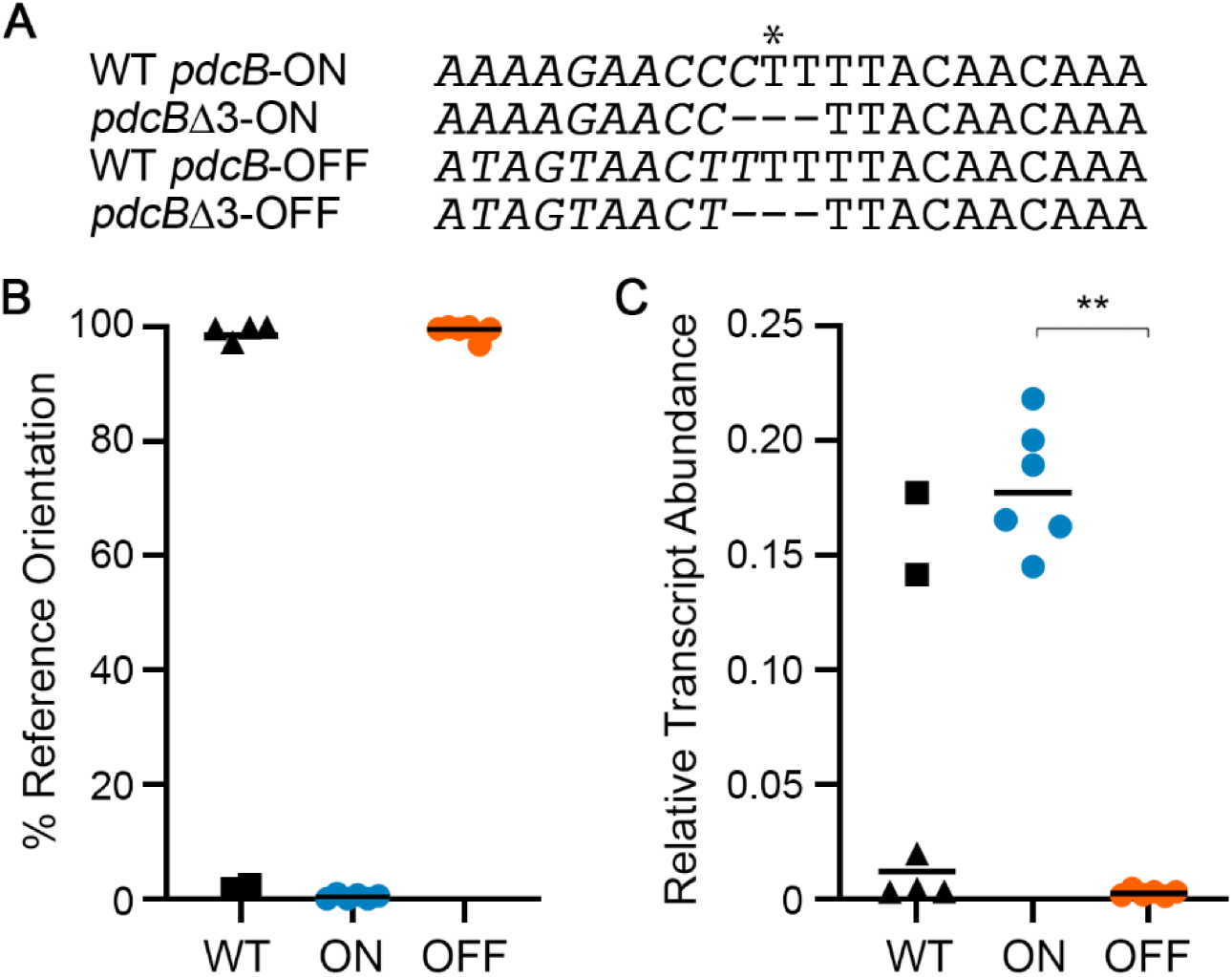
Mutation of the right inverted repeat results in phase-locked strains. (A) Three nucleotides in the RIR were deleted to create strains with the invertible element locked in the ON and OFF orientations. Asterisk indicates the site of recombination (19). Italics indicate a portion of the *pdcB* switch sequence that undergoes inversion. (B) Orientation-specific quantitative PCR was performed using genomic DNA from R20291 (WT), *pdcB*Δ3-ON, and *pdcB*Δ3-OFF. Data are expressed as the percentage of the invertible element in orientation present in the reference genome (published/OFF). (C) Quantitative reverse transcription PCR was performed using cDNA derived from R20291 (WT), *pdcB*Δ3-ON, and *pdcB*Δ3-OFF. Data are expressed as the ratio of the *pdcB* transcript abundance to that of the *rpoC* reference gene. (B, C) Symbols represent values from independent samples; bars indicate the median. For WT, triangles and squares distinguish isolates with reference and inverse sequence orientations, respectively. Means and standard deviations are shown. **p < 0.01 by Kruskal-Wallis test and Dunn’s post-test.

Due to the heterogeneity of *pdcB* switch orientation, we created mutants in which the *pdcB* switch was locked in the published or inverted orientation to further examine the effect of the switch orientation in *pdcB* expression and the role of *pdcB* in *C. difficile* physiology. Phase-locked mutants have previously been generated by inactivating the *recV* gene encoding the recombinase required for inversion of the flagellar and *cmr* switches (10-12), however the site-specific recombinase required for *pdcB* switch inversion has not been identified and is not RecV (19). Sekulovic et al previously identified the nucleotide in the inverted repeats where recombination occurs for each switch (19). Deleting this nucleotide and a nucleotide on each side was recently demonstrated to prevent site-specific recombination and inversion of the flagellar and *cmr* switches (46, 47). We used this strategy to create strains with the equivalent 3-nucleotide deletion in the right inverted repeat (RIR) of the *pdcB* switch, one in which the switch is in the reference genome orientation, and the other with the switch in the inverse orientation (Fig 2A). Whole genome sequencing confirmed the intended mutations and integrity of the genomes compared to the R20291 parent. We then applied OS-qPCR to confirm that the *pdcB* switch in these strains is locked in the desired orientations. As anticipated, the mutant with the *pdcB* switch in the presumed OFF orientation contained the switch in almost exclusively in the reference genome orientation (99.27% ± 1.21% reference), and we named this mutant *pdcB*Δ3-OFF (Fig 2B). The mutant with the switch in the presumed ON orientation had the inverted sequence (0.38% ± 0.42% reference), and we named this mutant *pdcB*Δ3-ON (Fig 2B). Both *pdcB*Δ3 mutants expressed *pdcB* consistent with their ON and OFF assignments, with levels equivalent to the naturally arising ON and OFF variants of wild-type R20291 (Fig 2C). Together these findings indicate that *pdcB* switch inversion mediates the phase variable expression of *pdcB*, and that the switch orientation in the R20291 reference genome corresponds to the OFF orientation, and the inverse sequence corresponds to the ON state.

### The *pdcB* switch contains an invertible promoter

In many examples of phase variation by site-specific DNA recombination, the invertible element contains a promoter that, when properly oriented, promotes transcription of the adjacent gene(s) leading to the phase ON state; in the other orientation, the promoter is directed away from the gene(s) resulting in the phase OFF state (14). To determine if a promoter is present in the *pdcB* switch, we generated plasmid-borne transcriptional reporters in which the *pdcB* switch is fused to the *phoZ* reporter gene encoding an alkaline phosphatase (AP) (Fig 3A). Fusions were made for the *pdcB* switch in both the OFF orientation (Cdi2-OFF::*phoZ*) and ON orientation (Cdi2-ON::*phoZ*). To prevent switch inversion during the experiments, the left inverted repeat (LIR) was excluded from the constructs. A promoterless fusion was included as a control (promoterless ::*phoZ*). Each of these plasmids was introduced into R20291, and AP activity was measured for the resulting strains. We found that when the *pdcB* switch was in the OFF orientation, AP activity was similar to the promoterless control; when in the ON orientation, AP activity increased 6-fold, indicating the presence of an active promoter in the ON, but not OFF, *pdcB* switch orientation (Fig 3A). To begin to determine the location of the putative promoter in the ON orientation, we made serial truncations from the 5’ end of the *pdcB* switch in the ON orientation (Cdi2-ONtrunc1::*phoZ* and Cdi2-ONtrunc2::*phoZ*). Removal of one-third of the *pdcB* ON switch sequence resulted in a 25% reduction in AP activity compared to the full-length construct, though the level of activity remained significantly higher than the Cdi2-OFF and promoterless reporters (Fig 3A). In contrast, removal of two-thirds of the *pdcB* ON switch sequence reduced AP activity to the levels of the promoterless control. Together these data indicate that a promoter is present near the center of the *pdcB* ON switch.

**Figure 3.**
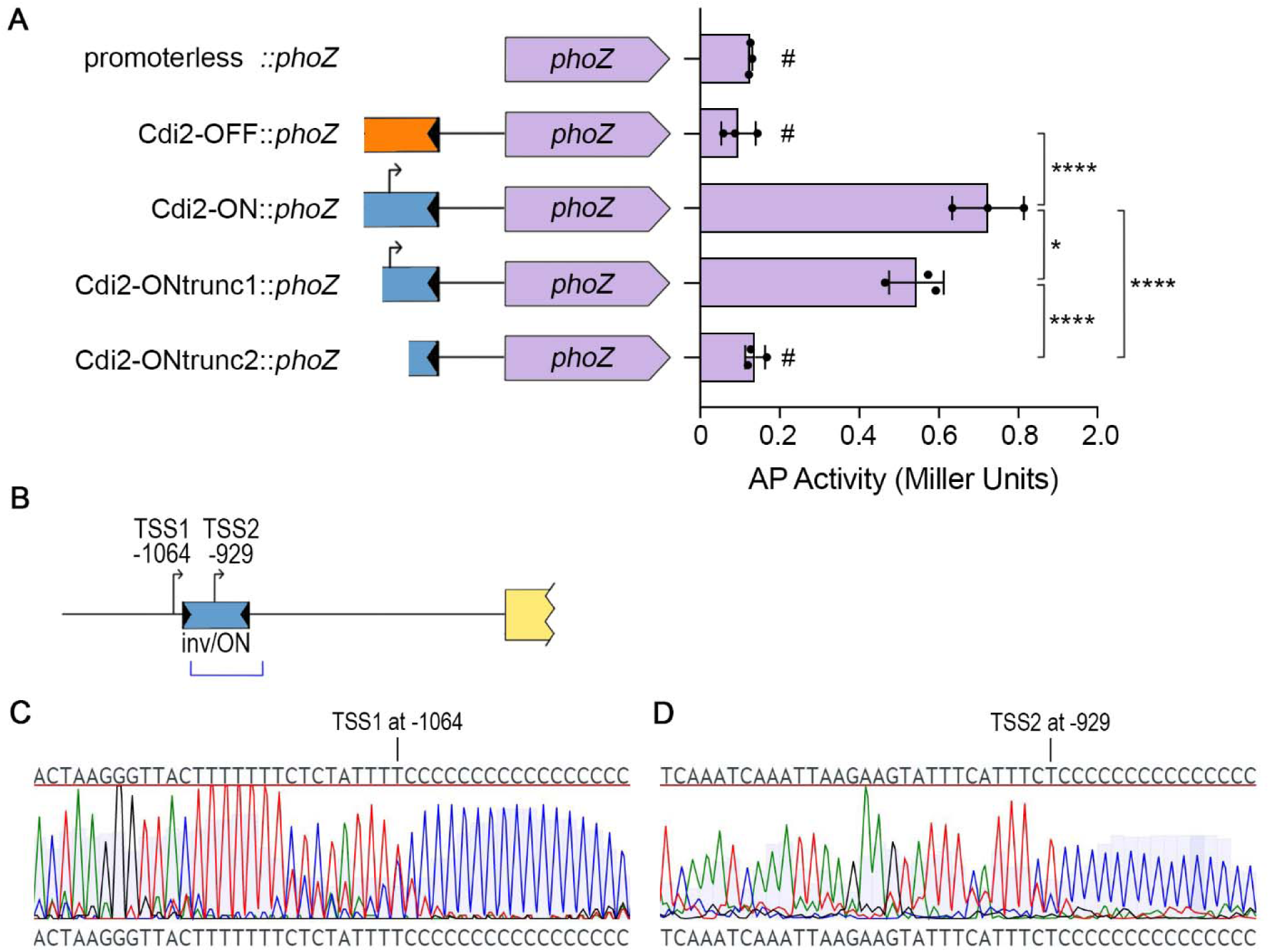
The *pdcB* switch contains an invertible promoter. (A) Alkaline phosphatase (AP) assay using *C. difficile* strains with plasmid-borne transcriptional fusions to *phoZ*. Promoterless *phoZ* was used as control. Means and standard deviations from 3 independent experiments are shown. *p < 0.05, ****p < 0.0001 by one-way ANOVA and Tukey’s post-test. No significant differences between strains marked #. (B) Diagram of the region upstream of *pdcB* with positions of TSS1 and TSS2 indicated with arrows. TSS2 appears in cDNA derived from the Cdi2 sequence in the inverse orientation (inv/ON). The blue bracket indicates the regions fused to *phoZ* in (A). (C,D) Chromatographs of the Sanger sequencing results obtained from the 5’ RACE experiment to identify transcriptional start sites (TSS) from (C) wild-type R20291 and (D) R20291 carrying the Cdi2-ON::*phoZ* reporter. TSS1 and TSS2 were identified as the first nucleotide adjacent to the poly-C tail added to the 5’ end of the cDNA created from the transcript of interest; the position relative to the translational start site is indicated.

To precisely map the position of the promoter present in the *pdcB* switch, we used 5’ Rapid Amplification of cDNA Ends (5’ RACE) to identify the transcriptional start site (TSS) in WT R20291. Because of the distance of the *pdcB* switch from the *pdcB* coding sequence, we used primers within 300 bp of the RIR. Using this strategy, we detected a TSS at position -1064 upstream of the annotated *pdcB* start codon (Fig 3B,C, Fig S1). This TSS is located upstream of the *pdcB* switch, 26 nucleotides 5’ of the LIR. We postulated that a promoter associated with this TSS might mask the signal from a promoter present in the *pdcB* switch, particularly since only a subpopulation of wild-type R20291 contains the switch in the ON orientation. To identify a TSS located specifically in the invertible element, we performed 5’ RACE using RNA extracted from*C. difficile* R20291 strains carrying the Cdi2-OFF::*phoZ* and Cdi2-ON::*phoZ* reporters that lack the LIR and therefore are unable to switch in *C. difficile*. For this experiment, we used a primer that anneals to *phoZ* to specifically identify any TSS from *phoZ* transcripts and not from the native locus. Using this approach, we identified a TSS from Cdi2-ON::*phoZ* corresponding to position -929 upstream of the predicted *pdcB* start codon (Fig 3B,D, Fig S1). No TSS was identified from Cdi-OFF::*phoZ*. These results indicate that at least two promoters regulate transcription of *pdcB*, one present when the switch is in the inverted/ON orientation and a second one present upstream of the *pdcB* switch (Fig 3B).

### *pdcB* expression affects intracellular c-di-GMP levels

*pdcB* encodes a protein with PAS, EAL, and GGDEF domains, and prior work showed that PdcB is a phosphodiesterase capable of hydrolyzing c-di-GMP (27). Phase variation of PdcB therefore is poised to modulate intracellular c-di-GMP levels. To test the effect of *pdcB* expression on c-di-GMP levels, we used a c-di-GMP riboswitch-based biosensor that allows measurements at the single cell level and in the bulk population. The biosensor, P_*gluD*_-PRS::mCherryOpt, is encoded on a multi-copy plasmid and consists of three elements: the heterologous *gluD* promoter that is not affected by c-di-GMP (31); the leader sequence of *pilA1* containing the Cdi-2-4 riboswitch that promotes *pilA1* gene expression and type IV pilus biosynthesis in response to c-di-GMP (31, 35); and the mCherryOpt gene encoding a red fluorescence protein codon-optimized for translation in *C. difficile* (48). An identical reporter with a mutation of a residue required for c-di-GMP binding was included as a negative control (P_*gluD*_-PRS^A70G^::mCherryOpt) (35, 36). The respective pPRS::mCherryOpt and pPRS^A70G^::mCherryOpt constructs were each introduced into WT, *pdcB*Δ3-OFF and *pdcB*Δ3-ON.

To measure c-di-GMP levels in these strains, we used a previously described assay to quantify red fluorescence normalized to the optical density of the culture (48, 49). For P_*gluD*_-PRS::mCherryOpt in the *pdcB*Δ3-ON background, we detected a significant 24% reduction in fluorescence compared to wild-type, consistent with increased production of PdcB and a corresponding reduction in c-di-GMP (Fig 4A, Fig S2). In the *pdcB*Δ3-OFF background, fluorescence was 65% higher than in the wild-type, indicating increased c-di-GMP as a result of decreased PdcB (Fig 4A, Fig S2). The equivalent strains bearing P_*gluD*_-PRS^A70G^::mCherryOpt, which cannot bind c-di-GMP, did not exhibit fluorescence above that of the R20291 control without the mCherryOpt plasmid (Fig S2). Thus, these reporters provided sensitive and specific detection of c-di-GMP *in vivo*. These results indicate that phase variation of PdcB significantly alters the global intracellular concentration of c-di-GMP in *C. difficile*.

**Figure 4.**
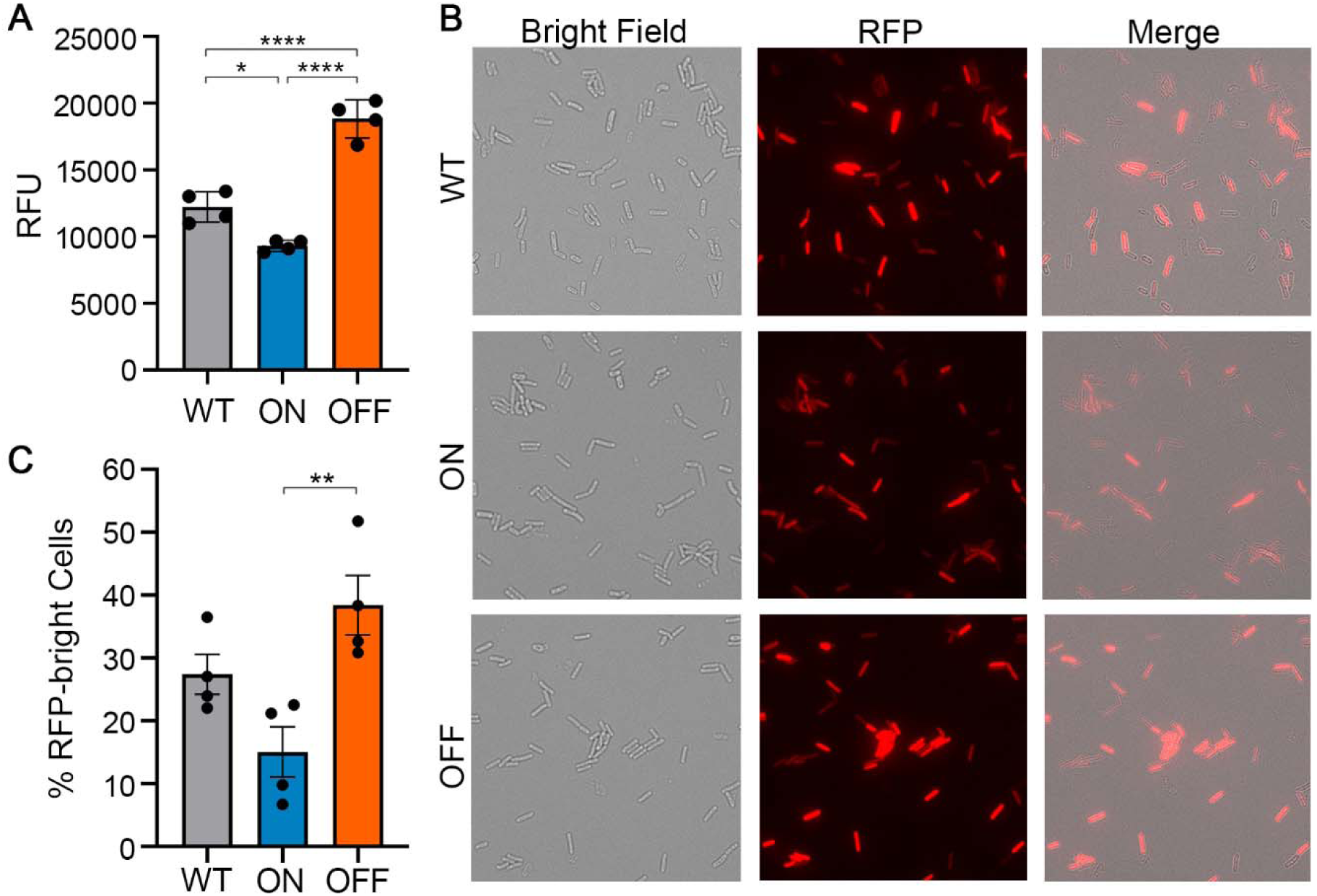
Phase variation of PdcB modulates c-di-GMP at the population and single-cell levels. (A) Measurement of c-di-GMP in vivo using the pPRS::mCherry reporter in R20291 (WT), *pdcB*Δ3-ON, and *pdcB*Δ3-OFF. Relative Fluorescence Units (RFU) calculated from arbitrary fluorescence units normalized to OD_600_. Kinetic analysis of fluorescence and data for the pPRS^A70G^::mCherry controls are in Fig S2. B) Representative micrographs showing heterogeneity in fluorescence in the above strains and variability in the number of bright red fluorescence protein-positive (RFP+) cells. (C) Quantification of bright RFP+ bacteria expressed as a percentage of total cells. Cells were counted and summed from three independent fields for four independent replicates of each strain. (A,C) Bars indicate means and standard deviations, with circles showing individual values. *p< 0.05, **p< 0.01, ****p<0.0001 by one-way ANOVA and Tukey’s post-test.

Particularly given heterogeneity of *pdcB* expression in wild-type R20291, the differences in c-di-GMP detected may arise from an increase in c-di-GMP in all cells, an increase in c-di-GMP in a subset of bacteria that increases the average of the bulk population, or both. To distinguish between these possibilities, we examined fluorescence of individual cells of wildtype, *pdcB*Δ3-ON, and *pdcB*Δ3-OFF bearing P_*gluD*_- PRS::mCherryOpt by microscopy. All three strains exhibited heterogeneity in the brightness of red fluorescence indicating variability in intracellular c-di-GMP across a population of *C. difficile*. Similar proportions of total bacteria exhibited some fluorescence: wildtype, 95.3% ± 2.1%; *pdcB*Δ3-ON, 93.4% ± 2.2%; and *pdcB*Δ3-OFF, 93.9% ± 0.2%) (Fig 4B). However, we observed differences in the proportions of bacteria with greater than baseline fluorescence, with significantly fewer RFP-bright bacteria in *pdcB*Δ3-ON compared to *pdcB*Δ3-OFF (Fig 4B,C). These results are consistent with the measurements of c-di-GMP in the bulk population and indicate that phase variation of PdcB impacts c-di-GMP level in a subpopulation of bacteria.

### Phase variation of PdcB modulates multiple c-di-GMP regulated processes in *C. difficile*

In *C. difficile*, c-di-GMP regulates multiple processes involved in motility and sessility. Because the *pdcB*Δ3 mutations significantly altered global c-di-GMP levels, we evaluated the effects of PdcB phase variation on representative c-di-GMP regulated phenotypes: swimming motility which is negatively regulated by c-di-GMP, and surface motility and biofilm formation which are positively regulated by c-di-GMP (33, 36, 41). In these assays, we compared wildtype, *pdcB*Δ3-ON, and *pdcB*Δ3-OFF; because *pdcB* expression can occur independently of the *pdcB* switch through the upstream promoter (Fig 3), we included a Δ*pdcB* control.

Swimming motility was assessed in BHIS-0.3% agar medium. The Δ*pdcB* and *pdcB*Δ3-OFF mutants were comparably motile, and both were significantly less motile than wildtype and *pdcB*Δ3-ON which exhibited similar motility to each other (Fig 5A). Surface motility was assayed on BHIS-1.8% agar medium. Here, the Δ*pdcB* and *pdcB*Δ3-OFF mutants exhibited comparable migration on the agar surface, and both showed significantly greater migration than wildtype and *pdcB*Δ3-ON, which again had similar phenotypes to each other (Fig 5B). These results are in accord with elevated c-di-GMP in the Δ*pdcB* and *pdcB*Δ3-OFF mutants resulting in inhibition of flagellar motility and augmentation of surface motility (12, 33, 47), and they imply that wildtype and *pdcB*Δ3-ON have similar c-di-GMP levels under these conditions.

**Figure 5.**
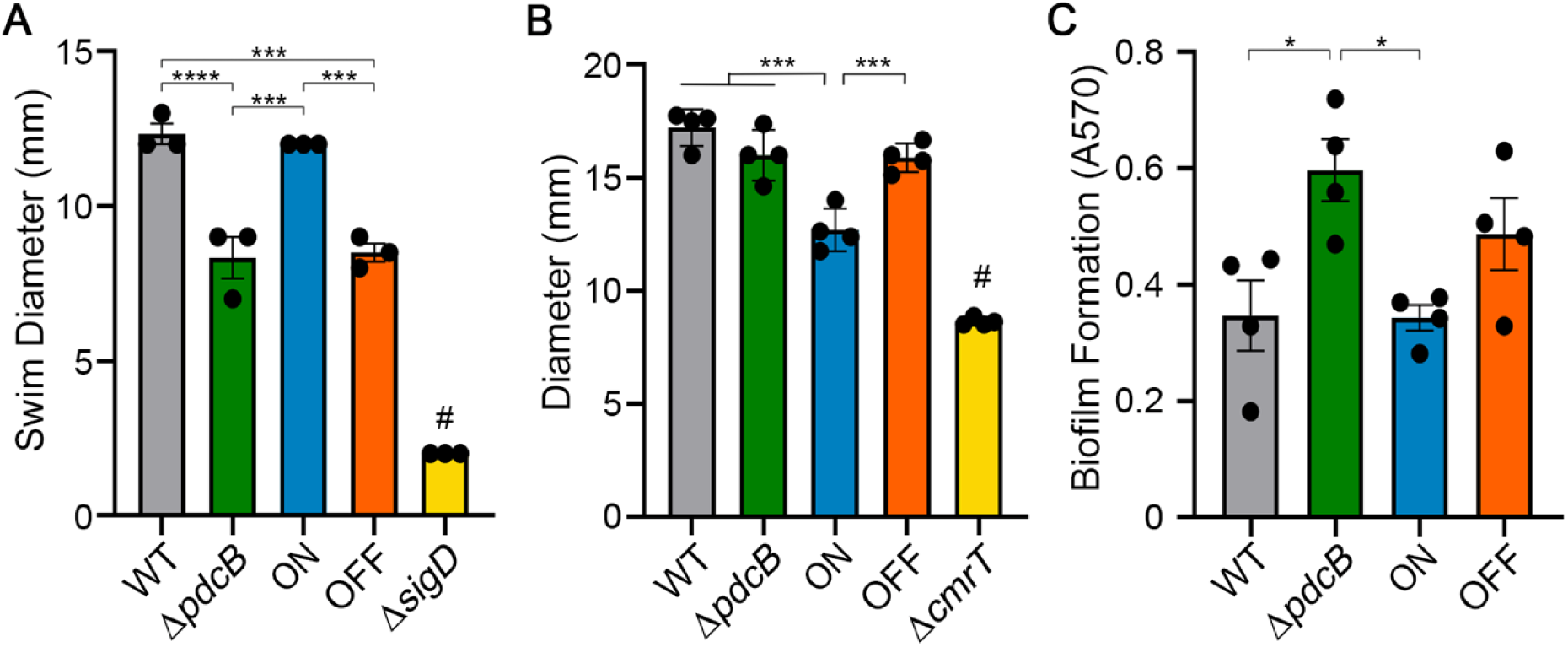
Expression of *pdcB* affects known c-di-GMP regulated processes. Phenotypic analysis of *C. difficile* R20291 (WT), Δ*pdcB, pdcB*Δ3-ON (ON), and *pdcB*Δ3-OFF (OFF). (A) Swimming motility in 0.5x BHIS-0.3% agar after 48 hours. A non-motile mutant (Δ*sigD*) served as a negative control. (B) Surface motility on BHIS-1.8% agar-1% glucose after 6 days. A Δ*cmrT* mutant served as a negative control. (C) Biofilm formation measured by crystal violet staining after 48 hours of growth in BHIS broth. (A-C) Means and error are shown. Symbols indicate values from independent samples. *p < 0.05, ***p < 0.001, ****p < 0.0001 by one-way ANOVA and Tukey’s post-test; #p< 0.05 compared to all other strains.

Biofilm formation was evaluated by culturing in BHIS medium in 96-well plates for 24 hours then quantifying adherent biofilm by crystal violet staining. In this assay, Δ*pdcB* and *pdcB*Δ3-OFF produced equivalent biofilm biomass, this time similar to wildtype (Fig 3C). *pdcB*Δ3-ON showed a modest but statistically significant 20% reduction in biofilm. These data support that PdcB hydrolysis of c-di-GMP inhibits biofilm development.

### Contribution of other phase variable factors

In this study we used mutants with fixed orientations of the *pdcB* switch, which prevents *pdcB* phase variation, to evaluate the role of PdcB in multiple c-di-GMP regulated processes. However, other phase variable factors could also influence these processes. The flagellar switch and *cmr* switch have been shown to impact swimming motility, surface migration, and biofilm formation (11, 12). In addition, a second c-di-GMP PDE gene, *pdcC*, is preceded by an invertible sequence (19). These factors might have phase varied in the mutants, leading to potential misattribution of function to PdcB. The functions of the remaining CDR20291_0963 and CDR20291_3417 loci are not yet known, and the upstream invertible sequences have not been shown to affect expression, so we cannot exclude a role in motility and biofilm formation. To address these possibilities, we first addressed the possible contribution of PdcC phase variation. Using *phoZ* reporter assays, we found that orientation of the *pdcC* switch modulates gene expression (Fig S3A). In addition, 5’ RACE detected a TSS when the *pdcC* switch is in the orientation that showed transcriptional activity (Fig S3B). Because *pdcC* exhibited phase variable expression, we determined the impact of PdcC on c-di-GMP regulated phenotypes of interest. Deletion of *pdcC* in R20291 had no effect on swimming motility or surface migration (Fig 6A, B). However, a Δ*pdcC* mutation led to a 2-fold increase in biofilm formation compared to wildtype, which is similar to the effect of the *pdcB* mutation on this phenotype (Fig 6C). Therefore, the increased biofilm of the Δ*pdcB* and *pdcB*Δ3-OFF mutants could also be attributed to PdcC.

**Figure 6.**
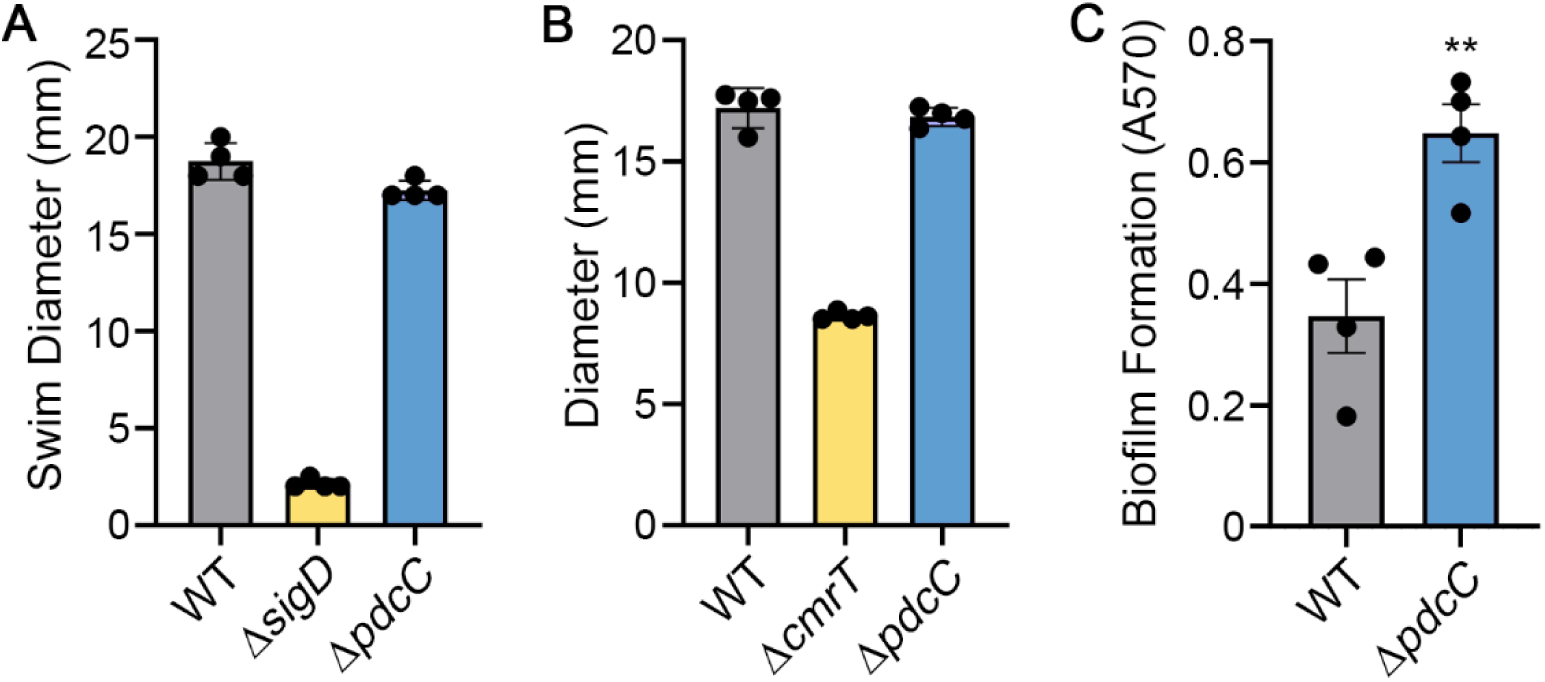
PdcC contributes to inhibition of biofilm formation, but not swimming or surface motility. (A) *S*wimming motility in 0.5x BHIS-0.3% agar after 48 hrs. A non-motile mutant (Δ*sigD*) served as a negative control. (B) Surface motility on BHIS-1.8% agar-1% glucose after 6 days. A Δ*cmrT* mutant served as a negative control. (C) Biofilm formation was measured by crystal violet staining after 48 hours of growth in BHIS broth. (A-C) Means and standard error are shown. Symbols indicate values from independent samples. **p < 0.01 by unpaired t-test.

We next determined the orientations of each of the other invertible sequences in the mutants used in this study using OS-qPCR. The proportions of the orientations of the *pdcB, pdcC*, flagellar (*flg*), *cmr, cwpV*, CDR20291_0963, and CDR20291_3417 invertible sequences were quantified for wildtype, Δ*pdcB, pdcB*Δ3-ON and *pdcB*Δ3-OFF grown in broth culture, expressed as the percentage in the orientation present in the R20291 reference genome. The *pdcB* switch was almost exclusively in the OFF, reference orientation, in all but the *pdcB*Δ3-ON as expected (Fig 7A). In all four strains, the *cmr, cwpV*, CDR20291_0963, and CDR20291_3417 sequences were in consistent orientations (Fig 7D-G), indicating that phenotypic changes observed in the *pdcB* mutants are not a result of phase variation of these sites. The *pdcC* and flagellar switches did differ among the strains, with wildtype and *pdcB*Δ3-OFF bearing these switches primarily in the reference orientation and Δ*pdcB* and *pdcB*Δ3-ON containing the inverse orientations (Fig 7B,C). These results indicate that inversion of the flagellar and *pdcC* switches occurred during the construction of Δ*pdcB* and *pdcB*Δ3-ON. However, the Δ*pdcB* and *pdcB*Δ3-OFF mutants exhibited the same motility and biofilm phenotypes despite having opposite orientations of the flagellar and *pdcC* switches, so these factors are unlikely to have contributed. We cannot dismiss the chance that phase variation occurred due to selective pressures present during growth in the assay conditions, but assessing the orientations of seven invertible sequences in multiple conditions with sufficient replicates for statistical power is imprudent without development of a higher throughput assay.

**Figure 7.**
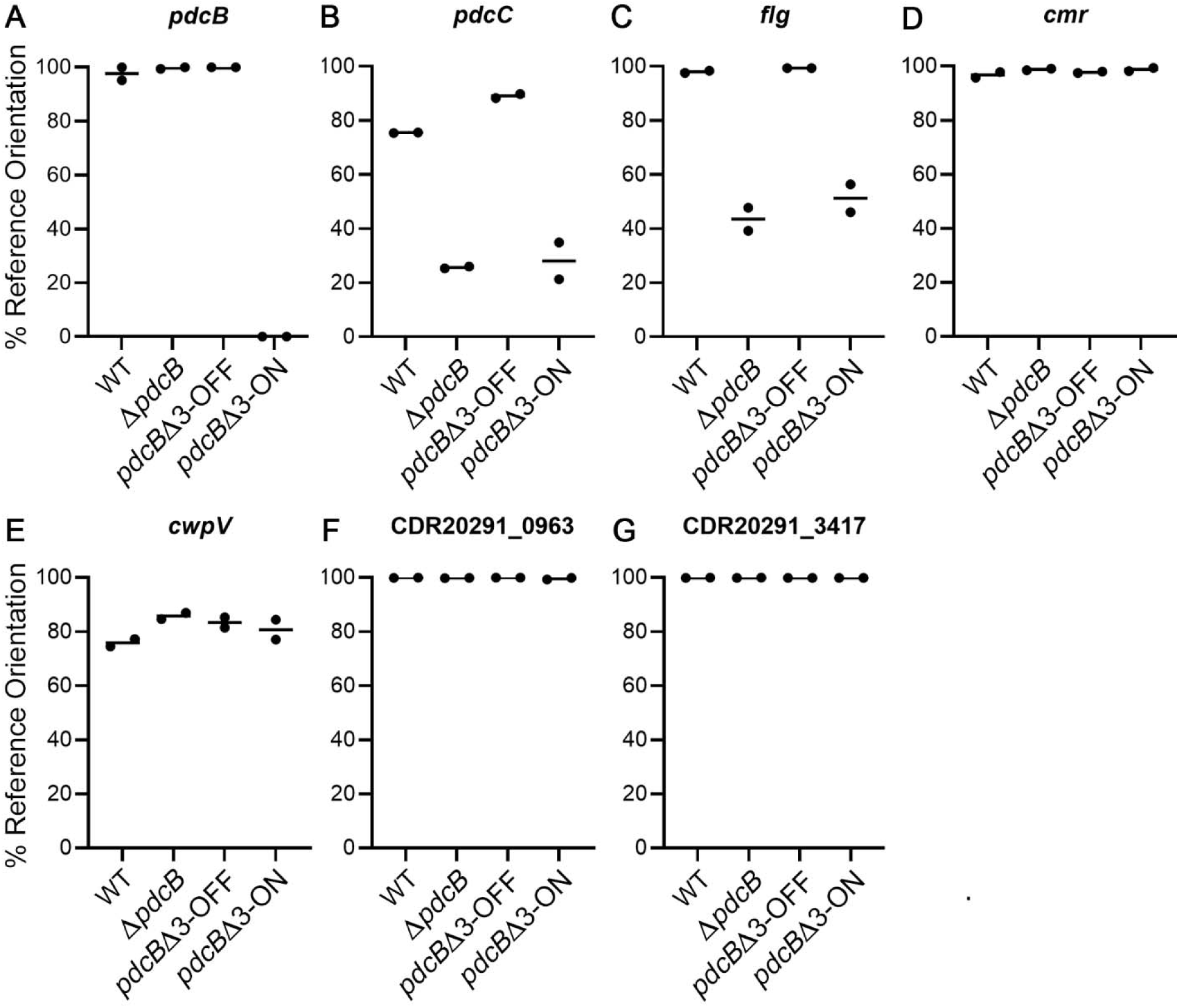
Orientations of the other invertible sequences do not correlate with the phenotypes of the *pdcB* mutants. Orientation-specific quantitative PCRs were performed using genomic DNA from wild-type R20291, Δ*pdcB, pdcB*Δ3-ON, and *pdcB*Δ3-OFF strains for the invertible elements *pdcB* (A), *pdcC* (B), *flg* (C), *cmr* (D), *cwpV* (E), CDR20291_0963 (F), and CDR20291_3417 (G). The data is presented as the percentage of the invertible element in the orientation of the reference R20291 genome sequence (Accession No. FN545816).

## DISCUSSION

In this study, we demonstrated that the *pdcB* switch controls phase variation of PdcB, modulating intracellular c-di-GMP levels and multiple known c-di-GMP regulated phenotypes in *C. difficile*. The effects of PdcB phase variation on c-di-GMP were apparent at the single cell and population levels. These findings indicate that phase variation of PdcB allows *C. difficile* to coordinate the control of multiple factors that play critical roles in motility, surface interactions, and biofilm formation, while diversifying these phenotypes in a population to ensure survival of environmental stresses.

Multiple *C. difficile* strains from different ribotypes showed heterogeneity in the orientation of the *pdcB* switch under standard laboratory conditions, suggesting that phase variation of PdcB is conserved and thus important for *C. difficile* fitness. However, the BAA1875 strain lacks the *pdcB* locus including the *pdcB* switch. Interestingly, this 078 ribotype strain also lacks flagellar genes. Given that flagellar genes and swimming motility are inhibited by c-di-GMP, BAA1875 may not experience the same selective pressures as strains that encode flagella. As a result, BAA1875, and perhaps other strains lacking flagella, may not need to modulate c-di-GMP to the same extent.

Analysis of paired DNA and RNA samples from single colonies showed that the ribotype 027 strain R20291 exhibits considerable heterogeneity in *pdcB* switch orientation and *pdcB* expression. Colonies with the *pdcB* switch in the reference genome orientation had lower *pdcB* transcript levels than colonies with the *pdcB* switch in inverse orientation, defining the OFF and ON states of the *pdcB* switch, respectively. These ON and OFF assignments are in agreement with the identification of a transcriptional start site (TSS2) within the *pdcB* switch when in the inverse/ON orientation but not in the published/OFF orientation. Further, an active promoter was detected using transcriptional reporters to the inverse/ON *pdcB* switch sequence, but not with a reporter to the published/OFF sequence. Interestingly, we identified an additional transcriptional start site (TSS1) upstream of the *pdcB* switch, which may control *pdcB* expression independent of phase variation. Whether and how these two putative promoters interact has not been determined. The region between the *pdcB* switch RIR and the annotated *pdcB* start codon is 840bp. No gene or regulatory RNA is predicted to be encoded in this region, and we did not detect a transcriptional start site in this region using 5’ RACE, suggesting that there is no promoter present between the *pdcB* switch and *pdcB*. It remains possible that this intervening sequence contains a promoter that is not active in the conditions tested, or that the long leader sequence of the mRNA forms a structure capable of further modulating expression.

The overall heterogeneity in *pdcB* switch orientation in wildtype R20291 impeded the ability to determine the phenotypic consequences of PdcB phase variation. Creating phase-locked strains by deleting 3 nucleotides from the RIR, rendering the *pdcB* switch unable to invert, circumvented this challenge and allowed us to determine the effect of PdcB phase variation. The phase-locked mutants differed significantly in *pdcB* expression. Single-cell analysis, using mCherryOpt under the control of a c-di-GMP-activated (*pilA1*) riboswitch as a reporter, revealed that the changes in c-di-GMP level were not uniform across the population. Instead, PdcB phase variation affected the number of bacteria exhibiting high fluorescence, with significantly more RFP-bright cells in *pdcB*Δ3-OFF (higher c-di-GMP) than in *pdcB*Δ3-ON (lower c-di-GMP). These differences may reflect the proportion of bacteria with c-di-GMP levels above the threshold needed for detection by the *pilA1* c-di-GMP riboswitch. The overall heterogeneity in fluorescence was not surprising since multiple DGC and PDE enzymes affect the c-di-GMP pool at any given time. Nonetheless, the orientation of the *pdcB* switch modulated the c-di-GMP level in the bulk population consistent with altered PdcB production and c-di-GMP hydrolysis; c-di-GMP was elevated in phase-locked OFF populations with lower *pdcB* expression and reduced in phase-locked ON populations overexpressing *pdcB*.

The phase-locked mutants showed changes in swimming motility, surface motility, and biofilm formation consistent with altered c-di-GMP – the locked OFF strain, with elevated c-di-GMP, exhibited reduced swimming motility, increased surface motility, and increased biofilm formation; the locked ON mutant showed the opposite phenotypic changes. The global effects on *C. difficile* physiology resulting from modulation of a single c-di-GMP PDE were not necessarily expected. Because *C. difficile* R20291 encodes 31 known or predicted c-di-GMP metabolic enzymes, there was a possibility of redundancy or compensation by another enzyme. Another PDE characterized in *C. difficile*, PdcA, only altered a subset of c-di-GMP regulated phenotypes (50). The broad impacts of PdcB suggest that this single PDE allows coordinated phase variation of multiple factors that are known or predicted to contribute to *C. difficile* virulence.

A challenge of studying phase variation in *C. difficile* is addressing the potential contributions of multiple switches to gene expression and physiology. For example, the flagellar and *cmr* switches impact swimming motility through distinct mechanisms.Further, a second phase variable c-di-GMP PDE, PdcC, is encoded in *C. difficile* R20291; deletion of *pdcC* reduced biofilm formation but did not affect swimming or surface motility. While preparing this manuscript, another report described motility, biofilm, and sporulation phenotypes of a *pdcB* mutant in another 027 ribotype strain, UK1 (51). However, that work did not account for phase variation of PdcC, or potential inversion of other switches. The colony morphology phenotype, for example, may have been attributable to phase variation of CmrRST. Heterogeneity in switch orientation may be masked by whole genome sequencing without specific efforts to detect inversions (19). In our study, we took steps to determine the orientations of all seven switches in the strains studied herein. While we cannot exclude the possibility that phase variation occurred during experimentation, we ensured that the starting switch orientations in each strain did not explain the observed phenotypes. Future work will determine whether the motility, biofilm, and other conditions select for specific phase variants and identify the potential selective pressures in these environments (52).

## MATERIALS AND METHODS

### Bacterial growth conditions

*C. difficile* strains were grown statically at 37°C in an anaerobic chamber with an anaerobic gas mix of 85% N_2_, 5% CO_2_, and 10% H_2_. Unless otherwise indicated, overnight cultures were grown in Tryptone Yeast (TY) broth, and cultures for experimentation were grown in Brain Heart Infusion medium (Becton Dickinson) supplemented with 0.5% yeast extract (Becton Dickinson) (BHIS). *Escherichia coli* strains were grown in Lucia Bertani (LB) medium (Fisher Scientific) with aeration at 37°C. Antibiotics were used at the following concentrations where indicated: chloramphenicol (Cm) 20 μg/mL, thiamphenicol (Tm) 10 μg/mL, kanamycin (Kan) 100 μg/mL, and ampicillin (Amp) 100 μg/mL.

### DNA manipulation and strain construction

To create the plasmids with *pdcB* (Cdi2) and *pdcC* (Cdi3) switch fusions to *phoZ*, the switch regions were PCR amplified from *C. difficile* R20291 genomic DNA with primers listed in Table S2. The PCR products were cloned into pMC123::*phoZ* (pRT1343) (47) and transformed into DH5α. Transformants were selected on LB-Cm agar and desired clones were identified by PCR and confirmed by sequencing. The plasmids were conjugated into *C. difficile* R20291 using *E. coli* HB101(pRK24) as the donor (53). Transconjugants were selected on BHIS-Kan-Tm agar (33); clones were screen by PCR with primers R837 and R839 to confirm the presence of the plasmid.

To create markerless deletions in the *C. difficile* genome, we used the toxin-antitoxin allele replacement system previously described (54). For deletion of *pdcB* (CDR20291_0685) and *pdcC* (CDR20291_1514), regions upstream and downstream of the locus to be deleted were amplified by PCR using primers named according to the convention locusF1/locusR1 for the upstream region and locus F2/locusR2 for the downstream region (Table S2). To mutate the RIR of *pdcB* switch, the upstream and downstream regions were amplified with primers introducing the desired mutation. Complementary overlapping sequences were added to the 5’ end of locusR1 and locusF2 primers to allow the products to be spliced together. To lock the *pdcB* switch in the ON orientation, primer sets R3082/R3092 and R3093/R3089 were used to amplify the upstream and downstream fragments, respectively. To lock the *pdcB* switch in the OFF orientation, R3082/R3090 and R3091/R3089 were used. The resulting upstream and downstream fragments were cloned into digested (*BamH*I and *Sac*I) pMSR0 vector using Gibson Assembly Master Mix (New England BioLabs). The desired clones were confirmed by PCR and sequencing of the inserts. These plasmids were conjugated into *C. difficile* strains using *E. coli* HB101(pRK24) as the donor. Allelic exchange was done as described (54). Candidate colonies were screened for the mutant allele using primers R3096/R3097 for Δ*pdcB*, R3146/R3147 for Δ*pdcC*, and R3096/R3138 for *pdcB*Δ3-ON and *pdcB*Δ3-OFF. The deletions were confirmed by sequencing. The resulting Δ*pdcB*, Δ*pdcC, pdcB*Δ3-ON and *pdcB*Δ3-OFF strains were confirmed to be isogenic with the R20291 parent by whole genome sequencing (Microbial Genome Sequencing Center, Pittsburgh, PA).

For the c-di-GMP reporter plasmid pP_*gluD*_-PRS::mCherryOpt and the pP_*gluD*_- PRS^A70G^::mCherryOpt negative control, the P_*gluD*_-PRS and P_*gluD*_-PRS^A70G^ fusions were amplified from previously generated *gusA* reporters containing these sequences, pRT942 and pRT943, respectively, by PCR using primers R3063 and R3064. The products were cloned by Gibson assembly into pDSW1728 digested with *Nhe*I and *Sac*I, replacing the existing P*tet* promoter with P_*gluD*_-PRS or P_*gluD*_-PRS^A70G^, to control expression of the mCherryOpt gene (48). Clones were identified by PCR and confirmed by sequencing of the inserts. The resulting pP_*gluD*_-PRS::mCherryOpt and pP_*gluD*_- PRS^A70G^::mCherryOpt fusions were introduced into R20291, *pdcB*Δ3-ON, and *pdcB*Δ3-OFF by conjugation via *E. coli* HB101(pRK24) as described previously, selecting on BHIS-Kan-Tm agar and confirming transconjugants by PCR (33, 53).

### Alkaline phosphatase assay

*C. difficile* colonies were inoculated into TY broth supplemented with Tm (TY-Tm) and grown overnight at 37°C. Cultures were diluted 1:30 into BHIS broth supplemented with Tm (BHIS-Tm) and grown to OD_600_ of approximately 1. Cultures were collected by centrifugation at 16,000 x g for 5 min. The supernatant was discarded and the pellets were saved at -80°C for 24 hrs. Pellets were thawed and the alkaline phosphatase (AP) assay was performed as previously described (55).

### 5’RACE

Overnight cultures of were diluted 1:30 in BHIS broth, with Tm as needed, and grown until OD_600_ of approximately 1. Cells were collected by centrifugation and stored in 1:1 ethanol:acetone. RNA was extracted and purified using the RNeasy Mini Kit (Qiagen) and treated with Turbo DNA-free™ Kit (Life Technologies). The 5’ RACE System for Rapid Amplification of cDNA Ends kit (Invitrogen™) was used to identify transcriptional start sites following the manufacturer’s protocol. Briefly, RNA was denatured, and cDNA was created by reverse transcription using primer GSP1. RNA was removed with the provided RNase mix and cDNA was purified using a S.N.A.P column. A homopolymeric tail was added to the 3’ end of the cDNA using terminal deoxynucleotidyl transferase TdT and deoxycytidine triphosphate (dCTP). Amplification of the cDNA was performed with a primer specific to the oligodC tail and a nested gene-specific primer (GSP2) (Table S2). PCR products were purified and sequenced to identify the TSS.

### Determination of switch orientation

Primers used are listed in Table S2. PCR with orientation-specific primers (OS-PCR) (Fig 1) was done using cell lysates as template (11, 19). Samples of overnight cultures were heated at 98°C for 10 minutes, then 1 μL was used as DNA template for PCR. Thermocycler conditions used were as follows: 98°C for 2 minutes, followed by 30 cycles of 98°C for 30 seconds, 54°C for 15 seconds, and 72°C for 30 seconds.

Quantitative OS-PCR (OS-qPCR) was performed as previously described (19). Standard OS-qPCR (Fig 2) used 100 ng genomic DNA as the template. When OS-qPCR was performed for all the switches (Fig 7), R20291, Δ*pdcB, pdcB*Δ3-OFF and *pdcB*Δ3-ON were cultured on BHIS agar plates; populations were obtained by collecting 100-150 colonies in 1 mL of BHIS broth. Genomic DNA was purified by phenol:chloroform:isopropanol (25:24:1) extraction and ethanol precipitation. of 20 μL that contained 100 ng DNA and primers at a final concentration of 100 nM each (Table S2). Reactions were run in a Lightcycler 96 system (Roche) with the following conditions: 98°C for 2 minutes followed by 40 cycles of 98°C for 30 seconds, 60°C for 60 seconds and 72°C for 30 seconds. A melting curve was performed as follows: 95°C for 10 seconds, 65°C for 60 seconds and 97°C for 1 second. The *rpoA* gene served as the reference. Data are expressed as the percentage of the indicated switch in the orientation present in the R20291 reference genome (Accession No. FN545816) (19).

### qRT-PCR

Overnight cultures grown in TY broth were diluted 1:30 in BHIS broth and grown to an OD_600_ of ∼ 1. Cells were collected by centrifugation, suspended in 1:1 ethanol-acetone, and stored at -80°C. RNA extraction, DNase treatment, and reverse transcription was performed as previously described (53). Real-time PCR reactions contained 4 ng of cDNA, primers at a final concentration of 500 nM each, and SYBR Green Real-Time qPCR reagents (Thermo Fisher). Reactions were run in a Lightcycler 96 system (Roche) as follows: 95°C for 10 minutes followed by 40 cycle amplification step of 95°C for 30 seconds, 55°C for 60 seconds and 72°C for 30 seconds. A melting curve was performed as follows: 95°C for 10 seconds, 65°C for 60 seconds and 97°C for 1 second. Data are expressed normalized to *rpoC*.

### RFP reporter assay

To quantify fluorescence produced by P_*gluD*_-PRS::mCherryOpt and P_*gluD*_- PRS^A70G^::mCherryOpt populations, fluorescence was monitored using a BioTek Synergy HT plate reader as described previously (48). Overnight cultures in BHIS-Tm (500 µl) were collected by centrifugation and suspended in 30 µl 1X DPBS. Washed cells (20 μL) was added to 180 µl of 1X DPBS in a flat-bottom clear 96-well microtiter plate (Corning 3370). The starting cell density (OD_600_) was recorded, then the contents of each well were transferred to a flat-bottom black 96-well microtiter plate (VWR 76221-764) for fluorescence readings. Fluorescence was recorded at 30-minute intervals using a Biotek HT plate reader (excitation, 590 nm; emission, 645 nm; sensitivity setting, 100) with a continuous slow shake for three hours.

### Microscopy

Cells were fixed to visualize population heterogeneity of reporter strains using a previously described protocol (48, 49). Briefly, overnight cultures of the P_*gluD*_- PRS::mCherryOpt and P_*gluD*_-PRS^A70G^::mCherryOpt reporter strains were grown overnight in BHIS-Tm. A 500 µl aliquot of culture strain was combined with 120 µl fixative (20 μl 1M NaPO_4_, pH 7.4; 100 µl 16% paraformaldehyde) and incubated 30 minutes at room temperature followed by 60 minutes on ice. Cells were washed with 1X DPBS before suspension in 500 µl 1X DPBS and incubation in the dark for two hours to allow for fluorophore maturation. Samples were applied to 1% agarose pads for microscopy using a Keyence BZ-X810 equipped with Chroma 49005-UF1 for RFP detection and a 60x oil immersion Nikon Plan Apo objective. Exposure and image capture settings were kept constant for all experiments. Four independent replicates of each strain were analyzed. Total cells, RFP-positive cells, and RFP-bright cells were counted from 5 images captured for each replicate. To identify RFP-bright cells, RFP channel images were processed identically using ImageJ by setting the threshold to 65.

### Phenotype Assays

Swimming: Overnight cultures (2 μL) were inoculated into 0.5X BHIS-0.3 % agar medium (11). Each plate contained all strains being compared. The diameter of growth was measured after 48 hours incubation at 37°C. Surface motility: Overnight cultures (5 μL) were spotted on the surface of BHIS-1.8% agar-1% glucose medium (12, 36). Each plate contained all strains being compared. Surface motility was determined on day 6 by measuring the largest diameter of each colony and a second, perpendicular measurement, then averaging the values (12). Biofilm formation: *O*vernight cultures were diluted 1:50 in BHIS and incubated at 37°C until they reached OD_600_ 0.5-0.6. The cultures were aliquoted (150 μL) into 96-well flat bottom plates (Corning) as five technical replicates. BHIS-only wells were included as controls. The plate was incubated for 48 hours, then removed from the anaerobic chamber for quantification of biofilm formation using crystal violet staining essentially as described previously (36, 37). Staining was measured by determining the absorbance at 570 nm. Technical replicate measurements were averaged for each biological replicate value.

## Acknowledgements

The authors declare no competing interests for this study. We thank former undergraduate research assistant Marlyn Anguelov for assistance with cloning. This work was supported by NIH awards R01-AI107029 and R01-AI143638 to R.T. L.R.R. was supported by a SPIRE Fellowship under NIH award K12-GM000678. The funding agency had no role in the design or execution of this study, the analysis of the data, or the decision to publish the results.

## SUPPLEMENTAL MATERIAL

**Table S1. Strains and plasmids used in this study.**

**Table S2. Oligonucleotides used in this study**.

**Figure S1. Mapping of transcriptional start sites in or near the *pdcB* switch**. Map of the transcriptional start sites (TSS) identified by 5’RACE in wild type R20291 and Cdi2-ON::*phoZ*. Depicted are the sequences of the *pdcB* switch in the inverted/ON orientation (red) and published/OFF orientation (blue). The sequences corresponding to the inverted repeats are in bold text. TSS identified are indicated in green highlight. Putative -10 and -35 sequences in the promoters are underlined.

**Figure S2. Kinetics of fluorescence development in PRS::mCherryOpt strains**. Fluorescence produced by wild-type R20291, *pdcB*Δ3-ON and *pdcB*Δ3-OFF strains carrying the pP_*gluD*_-PRS::mCherryOpt plasmid was quantified over a 3-hour time course during which the fluorophore matures. R20291 with no plasmid and strains carrying pP_*gluD*_-PRS^A70G^::mCherryOpt, which encodes a riboswitch that is blind to c-di-GMP, were used as controls. Data are expressed as fluorescence units normalized to optical density, shown as the means and standard deviations for four biological replicates.

**Figure S3. The orientation of the *pdcC* switch controls *pdcC* expression**. (A) Alkaline phosphatase (AP) assay using *C. difficile* strains with plasmid-borne transcriptional fusions to *phoZ:* the *pdcC* switch (Cdi3) in the published orientation (OFF), in the inverted orientation (ON), and two truncations of the 5’ end of the *pdcC* switch in the inverted orientation (ON). Promoterless *phoZ* was used as control. Means and standard deviations from 3 independent experiments are shown. **** p < 0.0001 by one-way ANOVA and Tukey’s post-test. (B) The transcriptional start site (TSS) identified by 5’RACE in R20291. The sequence of the *pdcC* switch in the inverted/ON orientation is in red and the published/OFF orientation is in blue. The sequences of the inverted repeats are in bold text. Green highlight indicates the TSS identified. Putative -10 and -35 sites are underlined.

