## Supplementary material for "Coordinated modulation of multiple processes through phase variation of a c-di-GMP phosphodiesterase in *Clostridioides difficile*": Table S1

**Table S1. Strains and plasmids used in this study**

| ***Clostridioides difficile* strains** | | | |
|---|---|---|---|
| **Lab Notation** | **Strain Name** | **Description** | **Reference** |
| RT273 | *C. difficile* R20291 | Ribotype 027 strain (Genbank Accession # FN545816) | (1) |
| RT1124 | *C. difficile* 630 | Ribotype 012 strain (Genbank Accession # AM180355) | (2) |
| RT1065 | *C. difficile* UK1 | Ribotype 027 strain | (3, 4) |
| RT1125 | *C. difficile* VPI10463 | Ribotype 003 strain | (5) |
| RT1357 | *C. difficile* ATCC BAA 1875 | Ribotype 078 strain | ATCC |
| RT1358 | *C. difficile* ATCC 43598 | Ribotype 017 strain | ATCC, (6) |
| RT1566 | *sigD* | R20291 *sigD*::*ermB* | (7) |
| RT2622 | Δ*pdcB* | R20291 Δ*pdcB* (in-frame deletion of CDR20291_0685) | This work |
| RT2797 | *pdcB*Δ3-ON | R20291 with *pdcB* switch locked in the ON orientation by deletion of 3 nucleotides in the RIR | This work |
| RT2796 | *pdcB*Δ3-OFF | R20291 with *pdcB* switch locked in the OFF orientation by deletion of 3 nucleotides in the RIR | This work |
| RT2816 | WT pP$_{gluD}$-PRS::mCherryOpt | R20291 pDSW1728::P$_{gluD}$-PRS::mCherryOpt; fluorescent c-di-GMP reporter with *gluD* promoter, *pilA1* 5' UTR containing c-di-GMP riboswitch, and codon-optimized mCherry | This work |
| RT2813 | WT pP$_{gluD}$-PRS$^{A70G}$::mCherryOpt | R20291 pDSW1728::P$_{gluD}$-PRS$^{A70G}$::mCherryOpt; mutation in riboswitch (A to G at position +70 in 5'UTR) rendering it blind to c-di-GMP | This work |
| RT2810 | *ΔpdcB* pP$_{gluD}$-PRS::mCherryOpt | R20291 Δ*pdcB* pDSW1728::P$_{gluD}$-PRS::mCherryOpt; fluorescent c-di-GMP reporter with *gluD* promoter, *pilA1* c-di-GMP riboswitch, and codon-optimized mCherry | This work |
| RT2814 | Δ*pdcB* pP$_{gluD}$-PRS$^{A70G}$::mCherryOpt | R20291 Δ*pdcB* pDSW1728::P$_{gluD}$-CdPRS$^{A70G}$::mCherryOpt; mutation in riboswitch (A to G at position +70 in 5'UTR) rendering reporter blind to c-di-GMP | This work |
| RT2812 | *pdcB*Δ3-ON pP$_{gluD}$-PRS::mCherryOpt | R20291 *pdcB*-ON pDSW1728::P$_{gluD}$-CdPRS::mCherryOpt; fluorescent c-di-GMP reporter with *gluD* promoter, *pilA1* c-di-GMP riboswitch, and codon-optimized mCherry | This work |
| RT2815 | *pdcB*Δ3-ON pP$_{gluD}$-PRS$^{A70G}$::mCherryOpt | R20291 *pdcB*-ON pDSW1728::P$_{gluD}$-CdPRS$^{A70G}$::mCherryOpt; mutation in riboswitch (A to G at position +70 in 5'UTR) rendering reporter blind to c-di-GMP | This work |
| RT2811 | *pdcB*Δ3-OFF pP$_{gluD}$-PRS::mCherryOpt | R20291 *pdcB*-OFF pDSW1728::P$_{gluD}$-CdPRS::mCherryOpt; fluorescent c-di-GMP reporter with *gluD* promoter, *pilA1* c-di-GMP riboswitch, and codon-optimized mCherry | This work |
| RT2817 | *pdcB*Δ3-OFF pP$_{gluD}$-PRS$^{A70G}$::mCherryOpt | R20291 *pdcB*-OFF pDSW1728::P$_{gluD}$-CdPRS$^{A70G}$::mCherryOpt; mutation in riboswitch (A to G at position +70 in 5'UTR) rendering it blind to c-di-GMP | This work |
| RT2195 | WT pMC123::*phoZ* | R20291 pMC123::*phoZ;* promoterless negative control for alkaline phosphatase assay | This work |
| RT2095 | WT pMC123::Cdi2-ONtrunc1::*phoZ* | R20291 pMC123::Cdi2-ONtrunc1::*phoZ;* Truncated version #1 of Cdi2 in the ON orientation fused to *phoZ* | This work |
| RT2096 | WT pMC123::Cdi2-ONtrunc2::*phoZ* | R20291 pMC123::Cdi2-ONtrunc2::*phoZ;* Truncated version #2 of Cdi2 in the ON orientation fused to *phoZ* | This work |
| RT2097 | WT pMC123::Cdi2-OFF::*phoZ* | R20291 pMC123::Cdi2-OFF::*phoZ*; Cdi2 in the OFF orientation fused to *phoZ* | This work |
| RT2098 | WT pMC123::Cdi2-ON::*phoZ* | R20291 pMC123::Cdi2-ON::*phoZ*; Cdi2 in the ON orientation fused to *phoZ* | This work |

| | | | |
|---|---|---|---|
| RT2058 | WT pMC123::Cdi3-ONtrunc1::*phoZ* | R20291 pMC123::Cdi3-ONtrunc1::*phoZ;* Truncated version #1 of Cdi3 in the ON orientation fused to *phoZ* | This work |
| RT2059 | WT pMC123::Cdi3-ONtrunc2::*phoZ* | R20291 pMC123::Cdi3-ONtrunc2::*phoZ;* Truncated version #2 of Cdi3 in the ON orientation fused to *phoZ* | This work |
| RT2457 | WT pMC123::Cdi3-OFF::*phoZ* | R20291 pMC123::Cdi3-OFF::*phoZ*; Cdi3 in the OFF orientation fused to *phoZ* | This work |
| RT2474 | WT pMC123::Cdi3-ON::*phoZ* | R20291 pMC123::Cdi3-ON::*phoZ*; Cdi3 in the ON orientation fused to *phoZ* | This work |
| RT2621 | Δ*pdcC* | R20291 Δ*pdcC* (in-frame deletion of CDR20291_1514) | This work |
| ***Escherichia coli* strains** | | | |
| **Lab Notation** | **Strain name** | **Description** | **Reference** |
| AC472 | DH5α | *E. coli* F- φ80*lacZ*ΔM15 Δ(*lacZY*A-*argF*)U169 *recA1 endA1 hsdR*17(rκ -, mκ+) *phoA supE44 thi*-1 *gyrA96 relA1* λ- *tonA* | Invitrogen, (8) |
| RT270 | HB101(pRK24) | *E. coli* used in conjugations with *C. difficile*, $Ap^R$ | (9) |
| **Plasmids** | | | |
| **Lab notation** | **Plasmid name** | **Description** | **Reference** |
| pRT2460 | pMSR0 | *E. coli*-*C. difficile* shuttle vector for toxin/anti-toxin mediated allelic exchange | (10) |
| pRT2563 | pMSR0::Δ*pdcB* | For in-frame deletion of *pdcB* (CDR20291_0685) | This work |
| pRT2795 | pMSR0::*pdcB*Δ3-*ON* | For mutation in RIR to lock *pdcB* switch ON | This work |
| pRT2794 | pMSR0::*pdcB*Δ3-*OFF* | For mutation in RIR to lock *pdcB* switch OFF | This work |
| pRT2766 | p$P_{gluD}$-PRS:: mCherryOpt | pDSW1728::$P_{gluD}$-PRS::mCherryOpt | This work |
| pRT2767 | p$P_{gluD}$-$PRS^{mut}$:: mCherryOpt | pDSW1728::$P_{gluD}$-$PRS^{mut}$::mCherryOpt | This work |
| pRT1343 | pMC123::*phoZ* | Promoterless vector to use as negative control for alkaline phosphatase assay | (11) |
| pRT2017 | pMC123::Cdi2-ONtrunc1::*phoZ* | Truncated version #1 of Cdi2 (*pdcB* switch) in the ON orientation fused to *phoZ* | This work |
| pRT2018 | pMC123::Cdi2-Ontrunc2::*phoZ* | Truncated version #2 of Cdi2 (*pdcB* switch) in the ON orientation fused to *phoZ* | This work |
| pRT2019 | pMC123::Cdi2-OFF::*phoZ* | Cdi2 (*pdcB* switch) in the OFF orientation fused to *phoZ* | This work |
| pRT2020 | pMC123::Cdi2-ON::*phoZ* | Cdi2 (*pdcB* switch) in the ON orientation fused to *phoZ* | This work |
| pRT1643 | pMC123::Cdi3-Ontrunc1::*phoZ* | Truncated version #1 of Cdi3 (*pdcC* switch) in the ON orientation fused to *phoZ* | This work |
| pRT1644 | pMC123::Cdi3-Ontrunc2::*phoZ* | Truncated version #2 of Cdi3 (*pdcC* switch) in the ON orientation fused to *phoZ* | This work |
| pRT1963 | pMC123::Cdi3-OFF::*phoZ* | Cdi3 (*pdcC* switch) in the OFF orientation fused to *phoZ* | This work |
| pRT1962 | pMC123::Cdi3-ON::*phoZ* | Cdi3 (*pdcC* switch) in the ON orientation fused to *phoZ* | This work |
| pRT2562 | pMSR0::□*pdcC* | For in-frame deletion of *pdcC* (CDR20291_1514) | This work |
| pRT942 | pMC123::$P_{gluD}$-PRS::*pilA1* | Used to amplify the $P_{gluD}$-PRS | |
| pRT943 | pMC123::$P_{gluD}$-$PRS^{A70G}$::*pilA1* | Used to amplify the $P_{gluD}$-$PRS^{A70G}$ | |
