## Supplementary material for "Coordinated modulation of multiple processes through phase variation of a c-di-GMP phosphodiesterase in *Clostridioides difficile*": Table S2

**Table S2. Oligonucleotides used in this study.**

| Lab notation | Primer name | Target[a,b] | Sequence (5'- 3') |
|---|---|---|---|
| R2215 | OS138 | Cdi1 PUB | CGCAATTATTTGTTTTTCATATGGATAAAATTGG |
| R2216 | OS139 | | GATTTTTATGTTAATGAATTGTTATAAAAAACATGG |
| R2213 | OS140 | Cdi1 INV | GGTAAGTTTGATTTTTATGTTAATGAATTG |
| R2214 | OS141 | | CAGTTTGTGCACTAGCTATGCCTGC |
| R2262 | OS101 | Cdi2 PUB | CATTTCTAAGAAATATCCTAACATAAAAACAAAA |
| R2263 | OS134 | | CGATTACACTACAGAATTAGAATGTCAATG |
| R2261 | OS100 | Cdi2 INV | GTTAAAAATTTAAGATATCTTTTCAGTATAATGGA |
| R2262 | OS101 | | CATTTCTAAGAAATATCCTAACATAAAAACAAAA |
| R2267 | OS106 | Cdi3 INV | GATTTGTCGAAACCATTGTAATAAGA |
| R2268 | OS107 | | CAATAGTTAAGACAATGAATATGCTACATTCT |
| R2268 | OS107 | Cdi3 PUB | CAATAGTTAAGACAATGAATATGCTACATTCT |
| R2269 | OS136 | | GTAAATTCCTCATAAAAATTTCCTCCCA |
| R2175 | qPCR_FlgSwit-ON | Cdi4 ON (PUB) | GTTTTCTTACCAAAGTGATACATTATTATATTAATG |
| R2177 | qPCR_FlgSwit-REV | | GCTATTGTCTGACTTCTTAAATTAGTTGCAT |
| R2176 | qPCR_FlgSwit-OFF | Cdi4 OFF (INV) | CATTAATATAATAATGTATCACTTTGGTAAGAAAAC |
| R2177 | qPCR_FlgSwit-REV | | GCTATTGTCTGACTTCTTAAATTAGTTGCAT |
| R2264 | OS104 | Cdi5 PUB | GTAAATTAAGATGTATTTCATTTCTCAAAAATATCCT |
| R2266 | OS135 | | GCTTTTATCGCAAGTTTGTTTTAAATGAC |
| R2264 | OS104 | Cdi5 INV | GTAAATTAAGATGTATTTCATTTCTCAAAAATATCCT |
| R2265 | OS105 | | GTAAAGTTTATAAAATCTGAAAAGCTCAAGA |
| R2271 | OS110 | Cdi6 PUB | CTAGCCAATAGACAAGTTTCTAGAAAAATA |
| R2272 | OS137 | | GAACAATTCTTGAATATTGTATTGAACATTAAGA |
| R2270 | OS109 | Cdi6 INV | GGAGATATATGGAGTTAGTGGTGCAA |
| R2271 | OS110 | | CTAGCCAATAGACAAGTTTCTAGAAAAATA |
| R2378 | OS196 | Cdi7 PUB | GTACAGAAGTTACCCAGAAGCTTGT |
| R2379 | OS197 | | TCCCCGCAATGGATGTTTTTTAATTCATC |
| R2378 | OS196 | Cdi7 INV | GTACAGAAGTTACCCAGAAGCTTGT |
| R2380 | OS198 | | TCCCAATTTAAATGTAGAGGTCATCAAT |
| R2273 | OS142 | *rpoA* | TCATTACCAGGTGTAGCAGTGAATGC |
| R2274 | OS143 | | TGATAGAGCATGGTCCTTGAGCTTCT |
| R3082 | CDR0685_F1 | Δ*pdcB* | CATTGATTTCTTTCAGTTTCGGATCCGTAACCCTTAGTTGTAAAAGGGTTC |
| R3083 | CDR0685_R1 | | CACTTTGATAGTTGGTCTAAACTTAAAGAGTTCGATTCTTCTC |
| R3084 | CDR0685_F2 | | CTCTTTAAGTTTAGACCAACTATCAAAGTGATGTACATAAAATGAG |
| R3085 | CDR0685_R2 | | GACGTCGACTCTAGAGGATCCCAGAACATTCCACGGTAAAATGG |
| R3086 | CDR0685IE_F1 | *pdcB*Δ3-OFF and *pdcB*Δ3-ON | CATTGATTTCTTTCAGTTTCGGATCCGTAATTGGTATACTTCCCTAGTTTACG |
| R3089 | CDR0685IE_R2 | | GACGTCGACTCTAGAGGATCCCATCTTTCCATATTTGAACATCATCC |
| R3090 | CDR0685IEpub_R1 | *pdcB*Δ3-OFF | GTAAAGTTACTATTTATTGAAAATTTAGATAC |
| R3091 | CDR0685IEpub_F2 | | CTAAATTTTCAATAAATAGTAACTTTACAAC |
| R3092 | CDR0685IEinv_R1 | *pdcB*Δ3-ON | GTAAGGTTCTTTTTTTTTATAATAAAATAGC |
| R3093 | CDR0685IEinv_F2 | | GCTATTTTATTATAAAAAAAAAGAACCTTAC |
| R3094 | phoZ-GSP1 | 5'RACE for TSS2 | CATCTTCTGGATAAGTG |
| R3095 | phoZ-GSP2 | | GCTTGCTGTCCGACCAAATAGGTATC |
| R3137 | GSP1_short-pdcB | | GGTACATTTTTAGTACATG |

| | | | |
|---|---|---|---|
| R3138 | GSP2_short-pdcB | 5'RACE for TSS1 | GAGTTGCTTACAGGGTATCTAGGATTGATG |
| R2419 | 0685inv_trunc1 | Cdi2::*phoZ* fusions | ATGCAGAATTCCTTAATTTGATTTGATATGTATATTTTTATAGC |
| R2420 | 0685inv_trunc2 | | ATGCAGAATTCGTTAAATTTATGAACATTTTTTGTTTTTATG |
| R2421 | 0685pub-LIR | | ATGCAGAATTCCTTTTTTTTTATAATAAAATAGCTATAAAAATATAC |
| R2422 | 0685inv-LIR | | ATGCAGAATTCCTATTTATTGAAAATTTAGATACTTTTCT |
| R2330 | 0685_pubR | | ATGCAGGATCCCGTACATAGTTTCCATTTGTTGTAAA |
| R3139 | R20291_0685qF | *pdcU* qRT-PCR | CCGGATGATGTTCAAATATGGAAAG |
| R3140 | R20291_0685qR | | TGGGTCATCCGACACATAAAC |
| R850 | rpoCqF | *rpoC* qRT-PCR | CTAGCTGCTCCTATGTCTCACATC |
| R851 | rpoCqR | | CCAGTCTCTCCTGGATCAACTA |
| R3063 | PgluD-CdPRS Gib | c-di-GMP biosensor | GAATTCTGCATCAAGCTAGCGAAAAGGAAATAATAGGAATATGGTAG |
| R3064 | PgluD-CdPRS Gib | | CTTTACTGCAGGAGCTCACTTACTTATAATATATCTGGTTAAAAATCTAAG |
| R3141 | CDR1514_F1 | Δ*pdcC* | CATTGATTTCTTTCAGTTTCGGATCCGTCTATCAAACTCGCTATAATATGTAGTG |
| R3142 | CDR1514_R1 | | CTTACCAAGTTGGACCTCATAAAAATTTCCTCCCATTAAAAC |
| R3143 | CDR1514_F2 | | GAAATTTTTATGAGGTCCAACTTGGTAAGAGGGTAATTAATCTG |
| R3144 | CDR1514_R2 | | GACGTCGACTCTAGAGGATCCGTCAACACCACCTATAGGTTCATC |
| R2218 | pdcCIE_pubF | Cdi3::*phoZ* fusions | ATGGAGCTCAATAAAATTTTTCAGACAATTCAAACAAAAATAATC |
| R2219 | pdcCIE-invF | | ATGGAGCTCTATCTACTTTAATTTTGTAAAATTTTCTGCTAC |
| R2220 | pdcCIE-R | | ATGGGTACCGCAAATAAGTACATCTCAAAAGTTTCC |
| R1977 | pdcC_IE_truncated1R | | TACGTGAATTCATTTATTAAAAGGTGATTATTTTTG |
| R1978 | pdcC_IE_truncated2R | | TACGTGAATTCGCATATTCATTGTCTTAACTATTG |
| R3096 | CDR0685_F0 | To screen for Δ*pdcB* allele | GCAAGAACCAATCAGTTACTTGAAG |
| R3097 | CDR0685_R0 | | GCTATGGAACATCCAGAAGAATATCC |
| R3146 | CDR1514_F0 | To screen for Δ*pdcC* allele | GCTTTATTACCAGCTATATGTGAAGAAC |
| R3147 | CDR1514_R0 | | CACTTATTAAGTGGGTAGCAATCAG |
| R837 | pUC19mcsF | To screen for pMC123 plasmid | TCTTCGCTATTACGCCAG |
| R839 | m13R | | AACAGCTATGACCATG |
| R3148 | GSP1_pdcC | 5'RACE for *pdcC* TSS | GACTCAAAGTTATTTATGG |
| R3149 | GSP2_pdcC | | GAGTTTGCATAAAGAATAGACTGGTGTTGTG |

[a] PUB indicates the intended template corresponds to that in the R20291 reference genome (FN585816); INV indicates the intended template has the inverted switch sequence

[b] Restriction sites are underlined
