## Supplemental Figures for "Coordinated modulation of multiple processes through phase variation of a c-di-GMP phosphodiesterase in *Clostridioides difficile*"

```
  -35                                   -10         TSS1
TTGAAATAGGGAAAAAATAGTGATATAATAAAAAAAATAGAGAAAAAAAGTAACCCT

TAGTTGTAAAAAAGTTACTATTTATTGAAAATTTAGATACTTTTCTAAAAATTATCT
   GTTGTAAAAGGGTTCTTTTTTTTTATAATAAAATAGCTATAAAAATATACATATC

                                                    -10        TSS2
ATGTTAATAGTTAAATTTATGAACATTTTTTGTTTTTATGTTAGGATATTTCTTAGA
AAATCAAATTAAGAAGTATTTCATTTCTAAGAAATATCCTAACATAAAAACAAAAAA

AATGAAATACTTCTTAATTTGATTTGATATGTATATTTTTATAGCTATTTTATTATA
TGTTCATAAATTTAACTATTAACATAGATAATTTTTAGAAAAGTATCTAAATTTTCA

AAAAAAAAGAACCCTTTTACAAC
ATAAATAGTAACTTTTTTACAAC
```

**Figure S1. Mapping of transcriptional start sites in or near the *pdcB* switch.** Map of the transcriptional start sites (TSS) identified by 5'RACE in wild type R20291 and Cdi2-ON::*phoZ*. Depicted are the sequences of the *pdcB* switch in the inverted/ON orientation (red) and published/OFF orientation (blue). The sequences corresponding to the inverted repeats are in bold text. TSS identified are indicated in green highlight. Putative -10 and -35 sequences in the promoters are underlined.

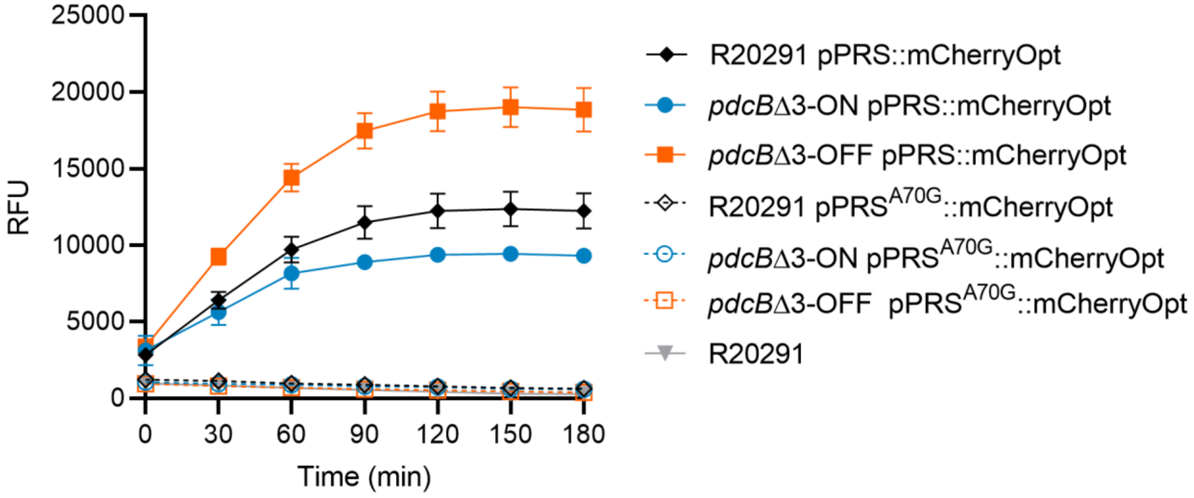


**Figure S2. Kinetics of P$_{gluD}$-PRS::mCherry fluorescence.** Fluorescence produced by wild-type R20291, *pdcB*Δ3-ON and *pdcB*Δ3-OFF strains carrying the pP$_{gluD}$-PRS::mCherryOpt plasmid was quantified over a 3-hour time course during which the fluorophore matures. R20291 with no plasmid and strains carrying pP$_{gluD}$-PRS$^{A70G}$::mCherryOpt, which encodes a riboswitch that is blind to c-di-GMP, were used as controls. Data are expressed as fluorescence units normalized to optical density, shown as the means and standard deviations for four biological replicates.

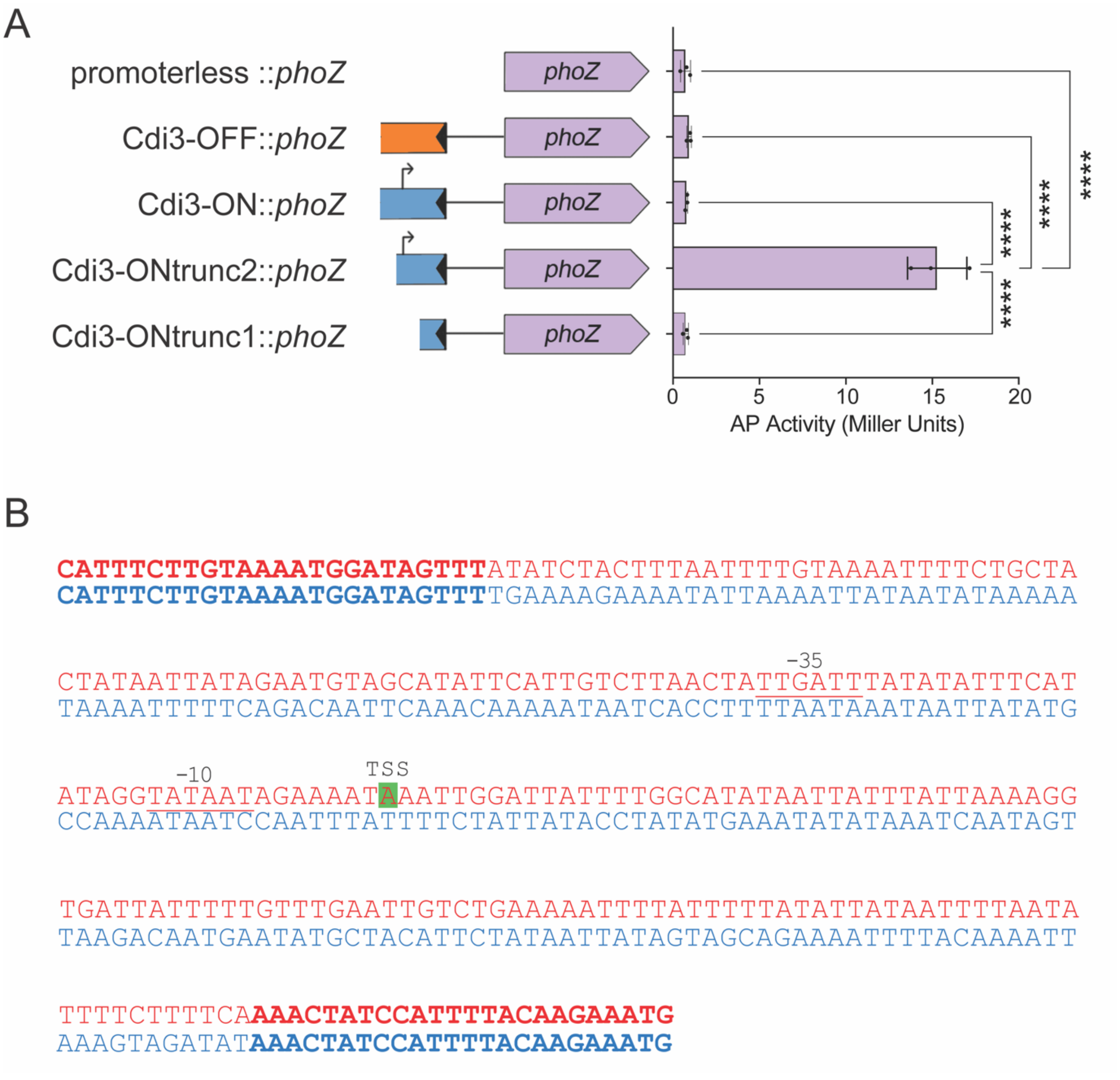


**Figure S3. The orientation of the *pdcC* switch controls *pdcC* expression.**
(A) Alkaline phosphatase (AP) assay using *C. difficile* strains with plasmid-borne transcriptional fusions to *phoZ:* the *pdcC* switch (Cdi3) in the published orientation (OFF), in the inverted orientation (ON), and two truncations of the 5' end of the *pdcC* switch in the inverted orientation (ON). Promoterless *phoZ* was used as control. Means and standard deviations from 3 independent experiments are shown. **** $p < 0.0001$ by one-way ANOVA and Tukey's post-test. (B) The transcriptional start site (TSS) identified by 5'RACE in R20291. The sequence of the *pdcC* switch in the inverted/ON orientation is in red and the published/OFF orientation is in blue. The sequences of the inverted repeats are in bold text. Green highlight indicates the TSS identified. Putative -10 and -35 sites are underlined.
